# MANIPULATING MITOCHONDRIAL REACTIVE OXYGEN SPECIES ALTERS SURVIVAL IN UNEXPECTED WAYS IN A *DROSOPHILA* MODEL OF NEURODEGENERATION

**DOI:** 10.1101/2024.03.25.586603

**Authors:** Andrew P. K. Wodrich, Brent T. Harris, Edward Giniger

**Affiliations:** National Institutes of Health, National Institute of Neurological Disorders and Stroke, Bethesda, MD; Georgetown University, Interdisciplinary Program in Neuroscience, Washington, DC; University of Kentucky, College of Medicine, Lexington, KY; Georgetown University, Department of Pathology, Washington, DC; Georgetown University, Department of Neurology, Washington, DC

**Keywords:** Cdk5, Mitochondria, Reactive oxygen species (ROS), Neurodegeneration

## Abstract

Reactive oxygen species (ROS) are associated with aging and neurodegeneration, but the significance of this association remains obscure. Here, using a *Drosophila* model of age-related neurodegeneration, we probe this relationship in the pathologically relevant tissue, the brain, by quantifying three specific mitochondrial ROS and manipulating these redox species pharmacologically. Our goal is to ask whether pathology-associated changes in redox state are detrimental for survival, whether they may be beneficial responses, or whether they are simply covariates of pathology that do not alter viability. We find, surprisingly, that increasing mitochondrial H_2_O_2_ correlates with improved survival. We also find evidence that drugs that alter the mitochondrial glutathione redox potential modulate survival primarily through the compensatory effects they induce rather than through their direct effects on the final mitochondrial glutathione redox potential *per se*. We also find that the response to treatment with a redox-altering drug varies dramatically depending on the age at which the drug is administered, the duration of the treatment, and the genotype of the individual receiving the drug. These data have important implications for the design and interpretation of studies investigating the effect of redox state on health and disease as well as on efforts to modify the redox state to achieve therapeutic goals.

## INTRODUCTION

Reactive oxygen species (ROS) are known to be produced in higher quantities with age and neurodegeneration, but the significance of these changes and their relationship to organismal health is unclear (Hou et al., 2019; Liu et al., 2017; Stefanatos & Sanz, 2018). Dating back to Harman’s free radical theory of aging, researchers have generally thought of ROS as detrimental byproducts of metabolism (Flohé, 2020; Harman, 1956). Indeed, an imbalance of redox homeostasis favoring high levels of ROS has been shown to be detrimental in some contexts (Shields et al., 2021). However, recent studies challenge the claim that ROS are invariably harmful (Lennicke & Cochemé, 2020; Ristow & Schmeisser, 2011; Shields et al., 2021). For example, inducing ROS production has been shown to extend lifespan in multiple species (Lee et al., 2010; Oka et al., 2015; Schroeder et al., 2013), while manipulations of antioxidant genes have surprisingly variable effects on organismal fitness despite altered levels of ROS (Page et al., 2010; Pérez et al., 2009; Ran et al., 2007; Van Raamsdonk & Hekimi, 2009). There remains no clear-cut answer to the question of whether ROS are detrimental by-products of metabolism or potentially beneficial signaling molecules, or whether either can be true, depending on the context (Wodrich et al., 2023). It also remains unclear whether the changes in redox state that occur in aging and neurodegeneration reflect a homeostatic response, contribute to dysfunction, or simply occur in parallel with the causative pathology. Moreover, it remains uncertain how aging and pathology modify the capacity to adapt to subsequent shifts in redox state.

Uncertainties remain, in part, due to the challenges of investigating, reporting, and discussing ROS. First, the umbrella term “ROS” is often used to discuss this entire class of molecules despite evidence that these molecules do not always act in concert (Lennicke & Cochemé, 2021; Sanz, 2016; Sies et al., 2022). Second, changes in a single reactive oxygen species have sometimes been viewed as reflective of a change in ROS generally (Lennicke & Cochemé, 2021; Murphy et al., 2022; Sanz, 2016). Third, certain methods of measuring ROS can be non-specific and, thus, difficult to unambiguously interpret due to molecular interactions with multiple ROS species or variable performance in differing pH environments, for example (Murphy et al., 2022; Sanz, 2016; Sies et al., 2022). Fourth, many studies that seek to investigate ROS do so by assessing proxy measures, such as lipid peroxidation, that correlate variably with changes in individual reactive oxygen species (Dotan et al., 2004; Murphy et al., 2022; Sanz, 2016).

Here, we use a simple *Drosophila* model of age-related neurodegeneration to investigate the above questions specifically as they relate to three different reactive oxygen species. Not only does this model have the inherent advantages of any *Drosophila* model of aging, namely the capacity for easy genetic manipulation and a short lifespan, but it also has the advantage that prior studies document the existence of transcriptional changes in mitochondrial redox-related processes that correlate with degeneration and survival (Shukla et al., 2022; Spurrier et al., 2018). This well-characterized model of age-related neurodegeneration relies upon altering the activity of the fly ortholog of a major human tau kinase, cyclin dependent kinase 5 (Cdk5; Pao & Tsai, 2021). Cdk5 is a phylogenetically conserved, noncanonical cyclin-dependent kinase that is associated with many age-related neurodegenerative diseases, including Alzheimer disease, Parkinson disease, and amyotrophic lateral sclerosis (Nguyen et al., 2001; Patrick et al., 1999; Qu et al., 2007; Smith et al., 2003). Cdk5 activity in mammals requires binding by one of its paralogous activating subunits, either p35 or p39, that are expressed only in postmitotic neurons, thereby limiting Cdk5 activity to that cell type (Pao & Tsai, 2021). In *Drosophila*, Cdk5 has only one activating subunit, the p35 ortholog Cdk5α, and knocking out Cdk5α eliminates Cdk5 activity altogether (Connell-Crowley et al., 2007; Kissler et al., 2009). Importantly, inactivating Cdk5 by knocking out Cdk5α (hereafter: Cdk5α-KO or KO) in flies causes several age-related neurodegenerative phenotypes such as shortened lifespan, axonal degeneration, impaired autophagy, motor impairment, altered innate immunity, and accelerated aging (Connell-Crowley et al., 2000, 2007; Howell et al., 2012; Kissler et al., 2009; Nandi et al., 2017; Shukla et al., 2019; Spurrier et al., 2018, 2019; Trunova & Giniger, 2012). Moreover, Cdk5α-KO flies demonstrate age-dependent degeneration of dopamine neurons and neurons in the mushroom body (MB), the region of the *Drosophila* central brain involved in learning and memory (Shukla et al., 2019; Spurrier et al., 2018; Trunova & Giniger, 2012).

Here we systematically measure three mitochondrial reactive oxygen species *in situ* in the brains of wild-type (hereafter: WT) and Cdk5α-KO flies at multiple ages, both with and without administration of drugs that have well-characterized effects on ROS. We then correlate the effects of the drugs on the levels of mitochondrial ROS with their effects on organismal survival to distinguish between beneficial, detrimental, or neutral effects on the animal. In particular, we ask whether pharmacological manipulations that either reverse or exacerbate the redox changes observed in Cdk5α-KO are beneficial or detrimental to the survival of these mutant flies. We find here that Cdk5α-KO flies have a reduced mitochondrial glutathione redox potential and lower mitochondrial H_2_O_2_ concentration across the lifespan than controls. Furthermore, we demonstrate that the lower mitochondrial H_2_O_2_ concentration is likely contributing to the reduced survival in these flies, and, indeed, that there is a general correlation between higher mitochondrial H_2_O_2_ concentration and improved survival. In contrast, drugs that modify the mitochondrial glutathione redox potential seem to modulate survival through the compensatory effects they induce and not by their effect on the final, net level of mitochondrial glutathione oxidation. Surprisingly, we do not uncover any clear evidence for a deleterious effect of the potent oxidizing agent, superoxide, though its level does change with age in WT flies. Finally, we observe multiple contexts where chronic exposure to various drugs that target redox pathways have effects that are very different to their effects after acute administration. This suggests that compensatory metabolic rewiring occurs in disease-relevant tissues in response to redox-altering drugs in a way that is both age- and genotype-dependent.

## RESULTS

### Baseline measurements of changes in three reactive oxygen species

To determine if the mitochondrial redox state is perturbed in aging and Cdk5 pathology, we used a series of reporters and dyes to assess three reactive oxygen species *in situ* in tissues that are known to be affected by Cdk5 pathology, namely the post-mitotic neurons of the brain and, specifically, the MB (Spurrier et al., 2018). We first used MitoSox-Red, a dihydroethidium-derived probe that is highly selective for superoxide within the mitochondria, to investigate changes in mitochondrial superoxide concentration within the whole brain (Robinson et al., 2006). At 10 days old, a young age when both WT and Cdk5α-KO flies are healthy, there is no difference in mitochondrial superoxide within the brain (Fig. 1). Over time, we observe an age-dependent increase in mitochondrial superoxide concentration in WT brains similar to reports elsewhere (Scialò et al., 2016); however, this age-dependent increase fails to occur in Cdk5α-KO (Fig. 1). To determine if aging or Cdk5 pathology alter the mitochondrial glutathione redox potential or H_2_O_2_ concentration, we used *201Y-GAL4* to express mitochondrially-localized redox biosensors (*UAS-mito-roGFP2-Grx1* or *UAS-mito-roGFP2-Orp1*, respectively) in the *Drosophila* MB, a region that undergoes age-dependent neurodegeneration (Albrecht et al., 2011; Spurrier et al., 2018). These biosensors have been used previously in other tissues in *Drosophila*, and we have validated that they also work reliably within the *Drosophila* MB (Supp. Fig. 1). In both 10 day-old and 30 day-old Cdk5α-KO adults, there is a reduction in the mitochondrial glutathione redox potential and lower mitochondrial H_2_O_2_ concentration relative to WT (Fig. 1; note that all groups were chronically fed a vehicle to permit comparisons with pharmacological challenge, described below; Supp. Table 1). It should be noted that there is a noticeable effect of the vehicle treatment specifically on the mitochondrial glutathione redox potential as compared to flies fed standard media without vehicle (Supp. Fig. 2A-C). Lastly, there do not appear to be global changes in ROS outside the mitochondria as we do not detect changes in the level of non-specific cytosolic ROS in the brain with age or Cdk5 pathology as measured by H_2_DCFDA (Supp. Fig. 2D).

**Figure 1.**
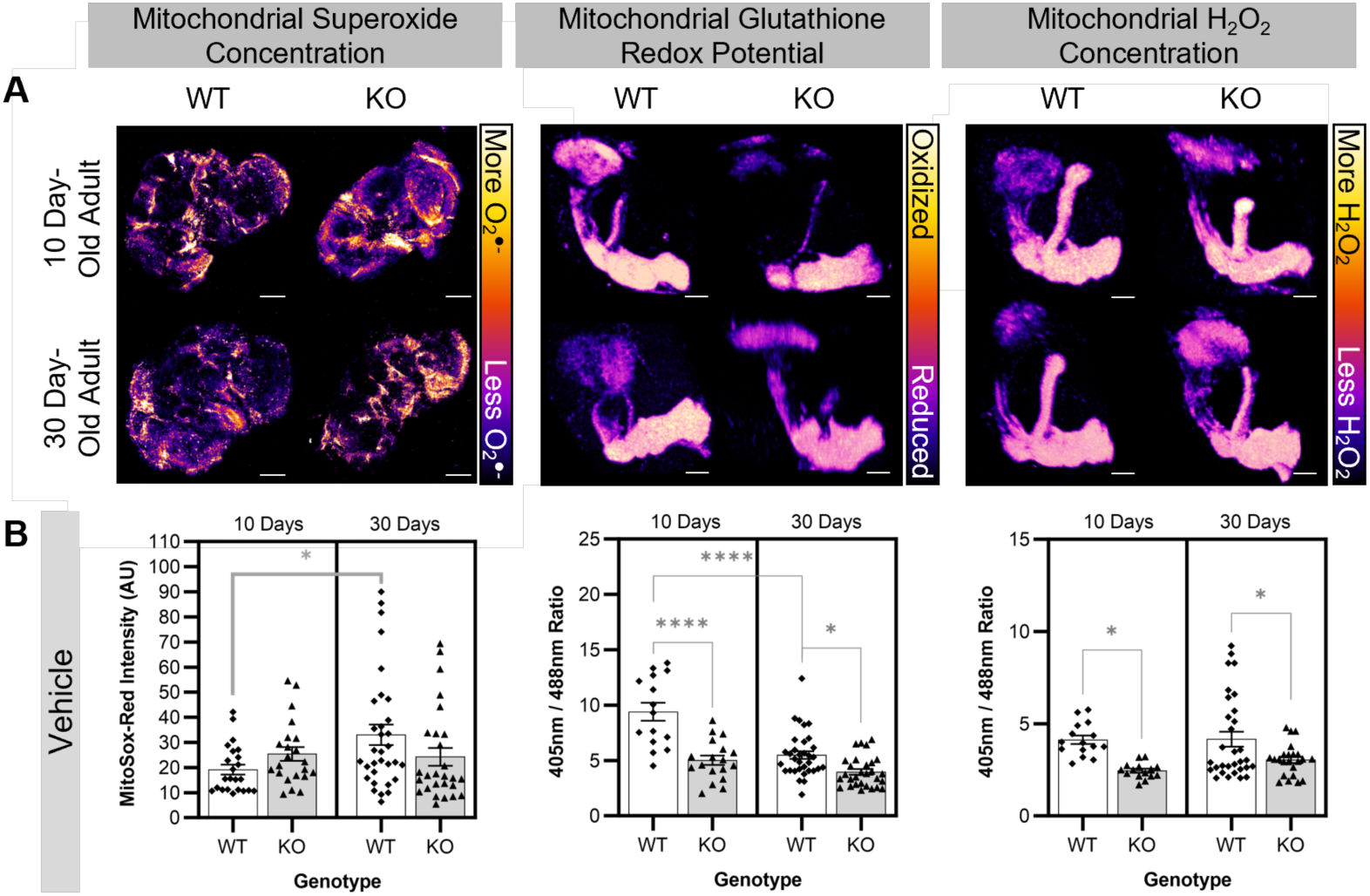
Baseline measurements of changes in three reactive oxygen species in WT and Cdk5α-KO. (A) Representative images of 10 day-old WT, 10 day-old Cdk5α-KO, 30 day-old WT, and 30 day-old Cdk5α-KO brains stained with MitoSox-Red, MBs expressing *201Y>mito-roGFP2-Grx1*, or MBs expressing *201Y>mito-roGFP2-Orp1* used to determine mitochondrial superoxide concentration, mitochondrial glutathione redox potential, or mitochondrial H_2_O_2_ concentration, respectively. For the mitochondrial superoxide concentration images, the scale bars represents 100 µm. For the mitochondrial glutathione redox potential and H_2_O_2_ concentration images, the scale bars represent 30 µm. For all images, the “Fire” look-up table was used with warmer colors (yellows and oranges) corresponding to higher ROS concentration or more oxidized glutathione redox potential and cooler colors (blues and purples) corresponding to lower ROS concentration or more reduced glutathione redox potential. (B) Quantification of mitochondrial superoxide concentration, mitochondrial glutathione redox potential, or mitochondrial H_2_O_2_ concentration in WT or Cdk5α-KO flies fed vehicle (DMSO or EtOH) chronically until aged 10 days old or 30 days old. Note that there are no significant differences based on the vehicle used. A higher MitoSox-Red intensity indicates a higher mitochondrial superoxide concentration. A larger 405nm/488nm ratio indicates a more oxidized mitochondrial glutathione redox potential (for *201Y>mito-roGFP2-Grx1*) or a higher mitochondrial H_2_O_2_ concentration (for *201Y>mito-roGFP2-Orp1*). Groups were compared using two-way ANOVA with Šidák’s multiple comparison test. *p* values: * < .05; ** < .01, *** < .001, **** < .0001.

**Table 1.**
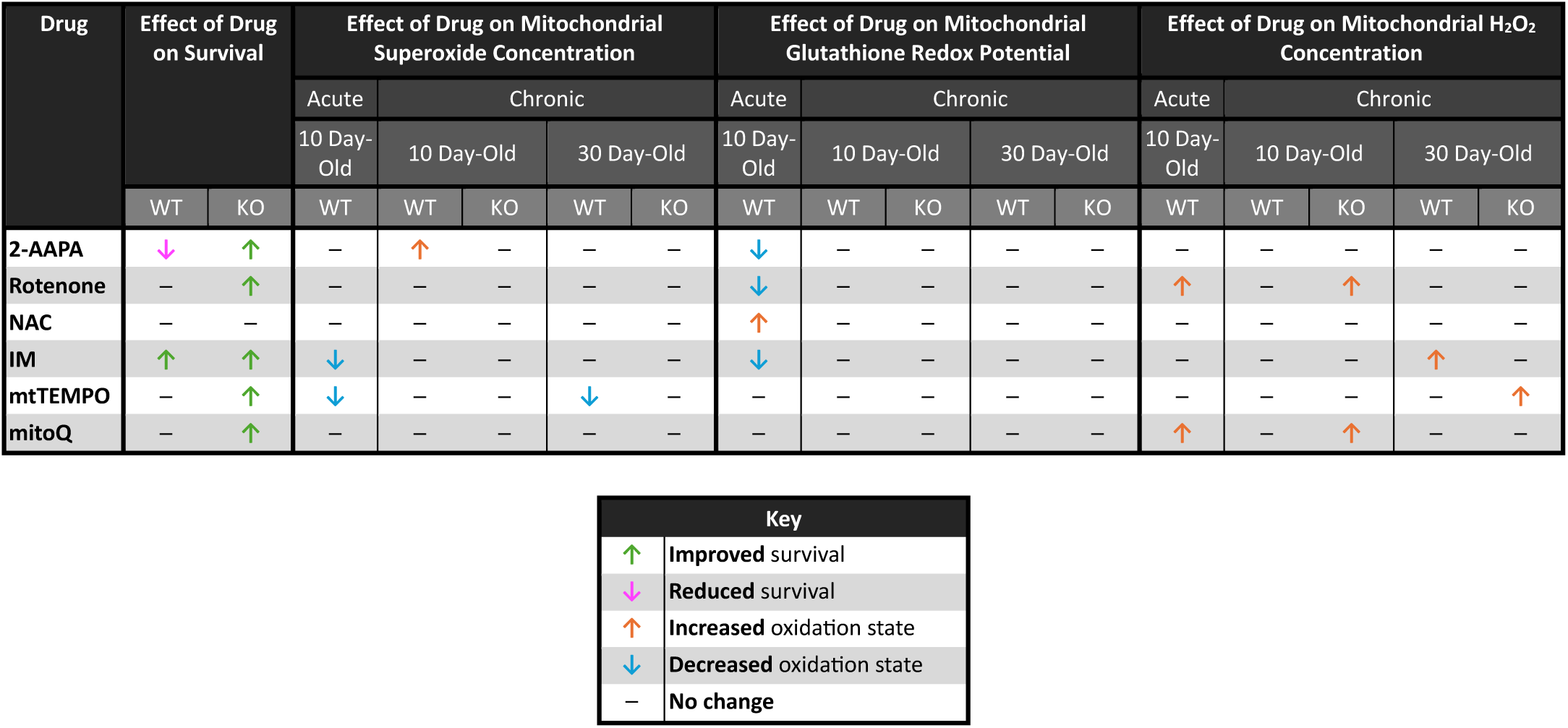
Summary of effects of acute and chronic drug administration on survival, mitochondrial superoxide concentration, mitochondrial glutathione redox potential, and mitochondrial H_2_O_2_ concentration in 10 day-old and 30 day-old WT and Cdk5α-KO flies. The observed drug effects are indicated by the green up arrow (drug improves survival), the magenta down arrow (drug reduces survival), the orange up arrow (drug increases oxidation state), the blue down arrow (drug decreases oxidation state), or a black bar (drug has no effect).

### Acute feeding of redox-altering drugs

To determine how the observed effects of aging and Cdk5α-KO on the three reactive oxygen species we have measured are related to organismal survival, we first needed the capability to reliably manipulate these redox parameters. We took a pharmacological approach to manipulating the mitochondrial redox state to have precise control over the timing and duration of manipulation and to avoid any potential disruption of development. We examined the literature to find drugs that have been reported to affect various redox parameters when fed to *Drosophila* (Supp. Table 2). Using dosages of these drugs intended to modestly affect the redox state without inducing toxicity, we performed a screen of 16 drugs predicted to affect one or more reactive oxygen species, based on their mechanisms of action as well as experimental evidence *in vitro* and *in vivo* (Supp. Tables 1, 2). To quantify the acute effects of these drugs, we fed them to 9 day-old WT adults for 24 hours before assessing the mitochondrial superoxide concentration, glutathione redox potential, and H_2_O_2_ concentration, as described above (Fig. 2A). We found 9 drugs that reliably alter the mitochondrial superoxide concentration, mitochondrial glutathione redox potential, or mitochondrial H_2_O_2_ concentration, or a combination thereof (Fig. 2B-D). We selected 6 of these drugs based on their observed effects on these redox parameters to use for subsequent experiments (Fig. 2E).

**Figure 2.**
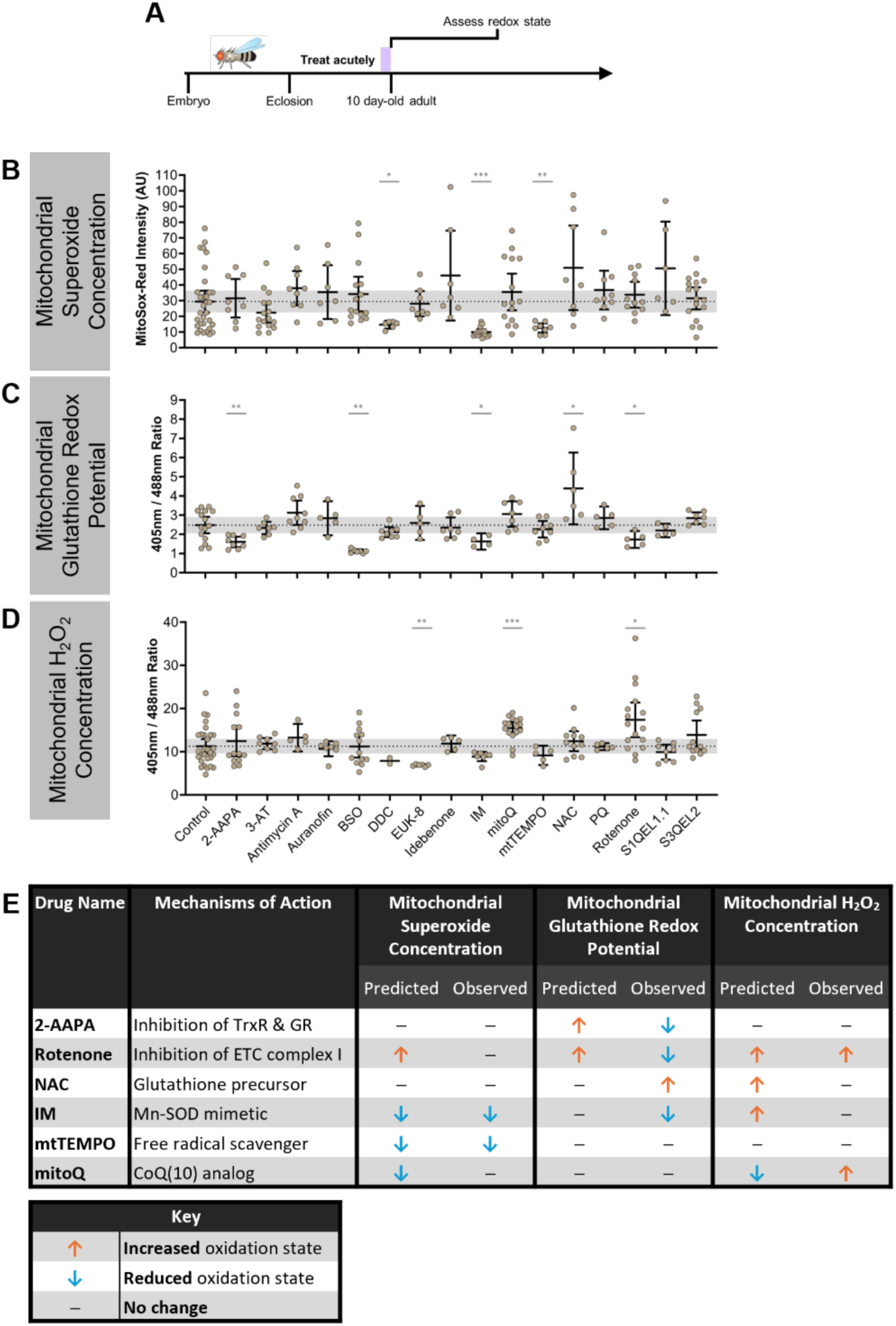
Acute feeding of redox-altering drugs. (A) Schematic of experimental design. WT flies were aged until 9 days old before feeding drugs acutely for 1 day. At 10 days old, flies were assessed for changes in mitochondrial redox parameters. (B-D) Quantification of mitochondrial superoxide concentration (B), mitochondrial glutathione redox potential (C), and mitochondrial H_2_O_2_ concentration (D) after acute feeding with 16 different drugs. A higher MitoSox-Red intensity indicates a higher mitochondrial superoxide concentration. A larger 405nm/488nm ratio indicates a more oxidized mitochondrial glutathione redox potential (for *201Y>mito-roGFP2-Grx1*) or a higher mitochondrial H_2_O_2_ concentration (for *201Y>mito-roGFP2-Orp1*). Error bars represent the 95% confidence intervals, and the gray shaded background represents the 95% confidence interval of controls. Differences between drug and control were compared using Kolmogorov-Smirnov tests. *p* values: * < .05; ** < .01, *** < .001, **** < .0001. (E) Summary table of predicted and observed effects of drugs selected for further analysis in this study. The drug effects are indicated by the orange up arrow (drug predicted to lead to an increased oxidation state), the blue down arrow (drug predicted to lead to a decreased oxidation state), or a black bar (drug predicted to led to no change in survival or oxidation state or a prediction cannot be made for the given drug).

### Chronic feeding of redox-altering drugs

To determine whether the observed age- and genotype-dependent changes in three reactive oxygen species contribute to organismal decline or are a homeostatic response to limit decline, we used the drugs identified above to alter the mitochondrial redox state in WT and Cdk5α-KO across the lifespan. We then assessed how rescuing or exacerbating the changes in ROS that we describe above affects survival in these flies. For chronic feeding, we fed drugs to WT or Cdk5α-KO flies from the day they eclosed as adults until they were 10 days old or 30 days old, at which point we assessed mitochondrial superoxide concentration, glutathione redox potential, and H_2_O_2_ concentration (Fig. 3A). We also assessed survival in response to drug treatment in both WT and Cdk5α-KO flies until they were 30 days old (Fig. 3A). Note that we were unable to reliably assess the effect of drug treatment on neuron cell loss in the MB since various drug treatments interfered with methods used to quantify neuron number. We found several changes in mitochondrial redox parameters and survival in response to redox-altering drug treatments. Specifically, we observe differences when we examine changes based on reactive oxygen species, drug, age, and genotype (Table 1). Here, we highlight salient observations.

**Figure 3.**
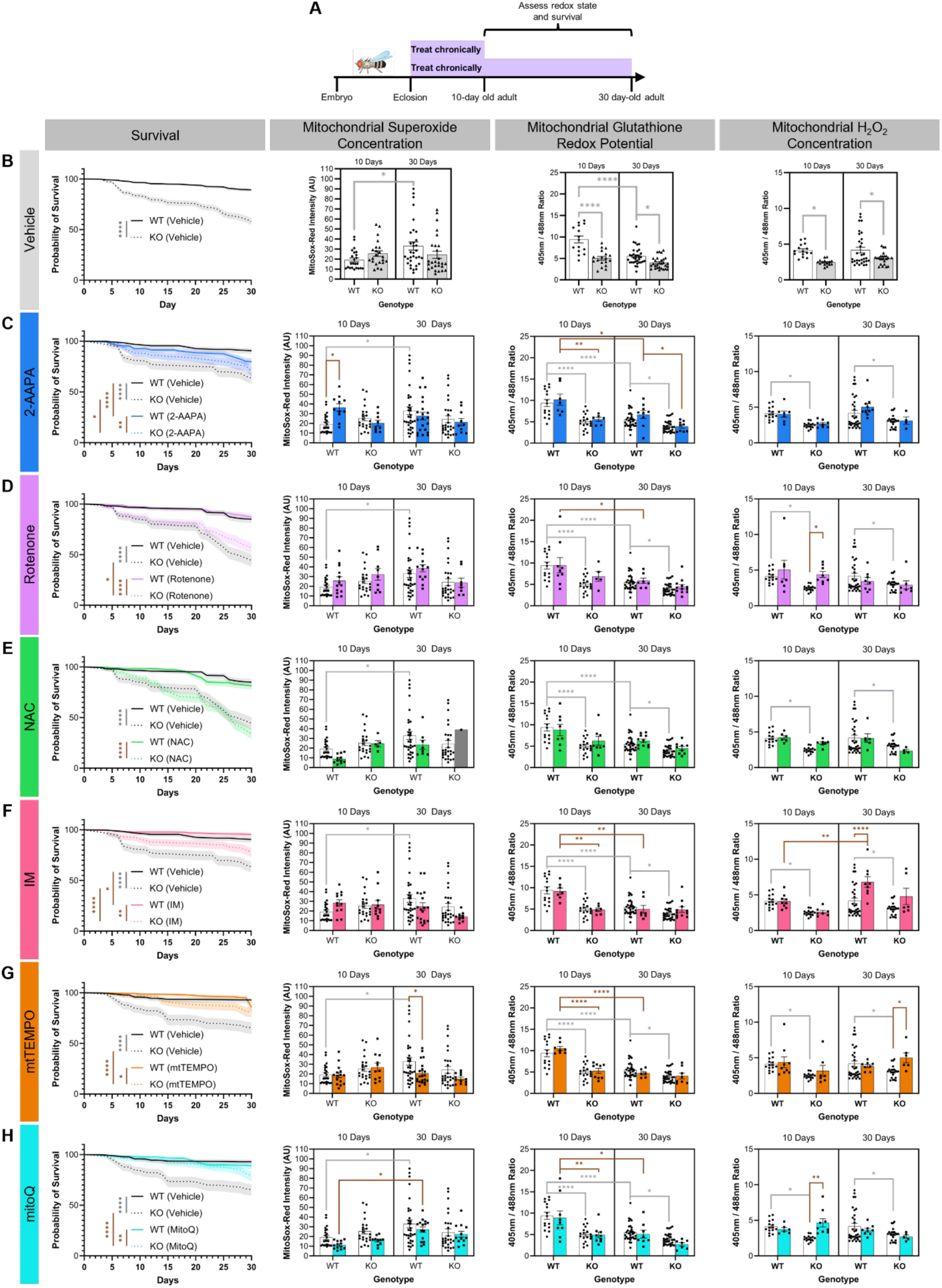
Chronic feeding of redox-altering drugs. (A) Schematic of experimental design. WT and Cdk5α-KO flies were chronically fed drugs from 0 days old until either 10 days old or 30 days old at which point flies were assessed for changes in mitochondrial redox parameters. Survival of all flies was monitored for 30 days. (B) Quantification of survival, mitochondrial superoxide concentration, mitochondrial glutathione redox potential, and mitochondrial H_2_O_2_ concentration in WT and Cdk5α-KO flies fed vehicle only. Note that the data in this panel is the same as shown in Figure 1B and is also used as the vehicle data in all the following panels of this Figure. A higher MitoSox-Red intensity indicates a higher mitochondrial superoxide concentration. A larger 405nm/488nm ratio indicates a more oxidized mitochondrial glutathione redox potential (for *201Y>mito-roGFP2-Grx1*) or a higher mitochondrial H_2_O_2_ concentration (for *201Y>mito-roGFP2-Orp1*). Shaded backgrounds in survival figures represent SEM. Differences in survival among groups were assessed by Mantel-Cox log-rank test with Bonferroni multiple comparison test. Differences in mitochondrial ROS among groups were assessed by two-way ANOVA with Bonferroni multiple comparison test. Light gray significance bars represent significant comparisons between vehicle groups and are presented in all the following panels. *p* values: * < .05; ** < .01, *** < .001, **** < .0001. (C-H) Quantification of survival, mitochondrial superoxide concentration, mitochondrial glutathione redox potential, and mitochondrial H_2_O_2_ concentration in WT and Cdk5α-KO flies fed 2-AAPA (C), rotenone (D), NAC (E), IM (F), mtTEMPO (G), or mitoQ (H). Note that vehicle controls are the same as shown in panel (B) and are shared for all drugs. Note that for the mitochondrial superoxide concentration measurement, N = 1 for 30 day-old Cdk5α-KO, and this is denoted by a gray bar (E). A higher MitoSox-Red intensity indicates a higher mitochondrial superoxide concentration. A larger 405nm/488nm ratio indicates a more oxidized mitochondrial glutathione redox potential (for *201Y>mito-roGFP2-Grx1*) or a higher mitochondrial H_2_O_2_ concentration (for *201Y>mito-roGFP2-Orp1*). Shaded backgrounds in survival figures represent SEM. Differences in survival among groups were assessed by Mantel-Cox log-rank test with Bonferroni multiple comparison test. Differences in mitochondrial ROS among groups were assessed by two-way ANOVA with Bonferroni multiple comparison test. Brown significance bars represent significant comparisons that include drug-treated groups. *p* values: * < .05; ** < .01, *** < .001, **** < .0001.

Looking across reactive oxygen species, we see numerous differences in how individual reactive oxygen species respond to various drug treatments. For example, none of the six drugs we assayed significantly alters the glutathione redox potential after 10 days or 30 days of chronic treatment as compared to vehicle-treated controls (Fig. 3). In contrast, rotenone and mitoQ both increase mitochondrial H_2_O_2_ concentration in 10 day-old Cdk5α-KO adults and this restoration of the mitochondrial H_2_O_2_ concentration towards the WT baseline correlates with a rescue of the Cdk5α-KO survival deficit, suggesting that the diminished mitochondrial H_2_O_2_ level in Cdk5α-KO may actually be detrimental to organismal survival (Fig 3). Consistent with this, increases in mitochondrial H_2_O_2_ in 30 day-old WT and Cdk5α-KO adults in response to IM and mtTEMPO, respectively, correlate with improvements in survival and further support a positive correlation between increased mitochondrial H_2_O_2_ concentration and improved survival (Fig. 3).

Looking across drugs, we observe that chronic administration of 2-AAPA has a discordant effect on survival between WT and Cdk5α-KO: WT survival is reduced by treatment with 2-AAPA while Cdk5α-KO survival is rescued (Fig. 3C). However, the only net change on mitochondrial ROS induced by chronic 2-AAPA is an increase in mitochondrial superoxide concentration in 10 day-old WT, which cannot by itself account for the differing survival effects in WT *vs.* Cdk5α-KO (Fig 3C). We also note that chronic administration of NAC, despite inducing oxidation of the mitochondrial glutathione redox potential after acute administration, has no effect on any of the mitochondrial redox parameters we examined after chronic administration, suggesting that drugs may exert different effects or be processed differently depending on the duration of drug administration (Fig 3E). Lastly, we see that chronic administration of either IM or mtTEMPO induces an age-dependent change in mitochondrial H_2_O_2_ concentration or superoxide concentration, respectively; that is, neither drug induces a change in mitochondrial redox parameters after 10 days of chronic administration, but they both induce effects after 30 days of chronic administration, suggesting that either the age of the animal or the duration of drug treatment is modulating its redox effect (Fig. 3F-G).

Several of our findings, including those described above, suggest that compensatory physiological responses to some drugs may have effects that alter or limit the direct effects of those agents, as determined by acute administration of these drugs to 10 day-old WT flies (Table 1). Thus, drugs administered acutely induce numerous significant effects on mitochondrial superoxide concentration, glutathione redox potential, and H_2_O_2_ concentration in 10 day-old WT flies. However, when these same drugs are administered chronically in either WT or Cdk5α-KO, many of these effects vanish. For the current analysis, we assume that the direct effect of each drug is similar in WT and Cdk5α-KO flies, but we cannot formally exclude the possibility that the acute effect of a drug may be altered in Cdk5α-KO.

As an example of compensation to chronic drug treatment, in WT flies, acute administration of 2-AAPA, rotenone, or IM reduces the mitochondrial glutathione redox potential, and NAC oxidizes it, but none of these affects the mitochondrial glutathione redox potential after chronic drug administration (Fig. 2, 3). The consequences of these examples of compensation are surprisingly varied, however, as IM improves survival, NAC and rotenone have no effect on survival, and 2-AAPA causes opposing survival effects in WT *vs.* Cdk5α-KO, as we will discuss in more detail below (Fig. 3). The ability to compensate for the effect of a drug, moreover, is apparently dependent on the genotype. We observe, for example, that Cdk5α-KO flies show changes in mitochondrial H_2_O_2_ concentration under chronic treatment with rotenone or mitoQ that match the direct effects of these drugs observed upon acute treatment; that is, they fail to compensate for the acute effects of these drugs on mitochondrial H_2_O_2_. In contrast, WT flies suppress completely the direct effect of these drugs in the same chronic treatment condition and show no change in final mitochondrial H_2_O_2_ concentration (Fig. 2, 3). This could imply that Cdk5α-KO impairs the homeostatic machinery that would normally compensate for the effects of drugs, or that the baseline redox state of Cdk5α-KO is intrinsically more sensitive to the effects of drugs. Acute administration of IM and mtTEMPO each causes a decrease in mitochondrial superoxide concentration while chronic administration of either drug for 10 days does not affect mitochondrial superoxide concentration in WT, suggesting that there is a compensatory response to these drugs that limits their effect on this redox parameter (Fig. 2, 3). Furthermore, the effect of the drug, the compensatory response to the drug, or a combination of these prevents the age-dependent increase in mitochondrial superoxide concentration that occurs without drug in WT (Fig. 3). However, the relationship between these effects on mitochondrial superoxide and survival cannot be determined from these data, as chronic administration of either IM or mtTEMPO also increases the mitochondrial H_2_O_2_ concentration which we have already found to be strongly associated with improved survival, as discussed above.

### Effects of redox-altering drug treatment duration vs. age at treatment assessment

Differences in the effects of a chronic drug treatment on 10 day-old *vs.* 30 day-old adults could be due to the absolute age of the animal at the time of assessment or the duration of drug treatment. Therefore, we tested whether the duration of drug treatment or the age at which the drug effect is assessed is more relevant for the response to the drug in 30 day-old adults. We compared mitochondrial superoxide levels in WT adults fed mtTEMPO or NAC either from eclosion to 10 days old, from eclosion to 30 days old, or for a 10-day period from 20 days old until 30 days old (Fig. 4A). Interestingly, we found evidence for each of the possible explanations, depending on the drug. For mtTEMPO, which shows a significant effect on mitochondrial superoxide concentration with treatment from 0 days to 30 days as compared to vehicle-treated controls, we find that feeding drug to WT flies just from 20 days old to 30 days old does not alter the mitochondrial superoxide concentration as compared to control, matching the effect of mtTEMPO feeding from eclosion until 10 days old (Fig. 4B). This suggests that the duration of treatment is more relevant for the effect of this drug than is the age of the animal at the time of assessment. In contrast, the results of feeding WT flies NAC from 20 days old until 30 days old match closely those of feeding NAC to WT flies from 0 days old until 30 days old and are significantly different from those of feeding NAC to WT flies from 0 days old until 10 days old, arguing that actual age at the time of assessment is more important for the observed effects of the drug rather than the duration of treatment (Fig. 4C). In total, this suggests that both treatment duration and age can contribute to the effect of a drug on aged flies, with the dominant factor depending on the specific pharmacologic agent.

**Figure 4.**
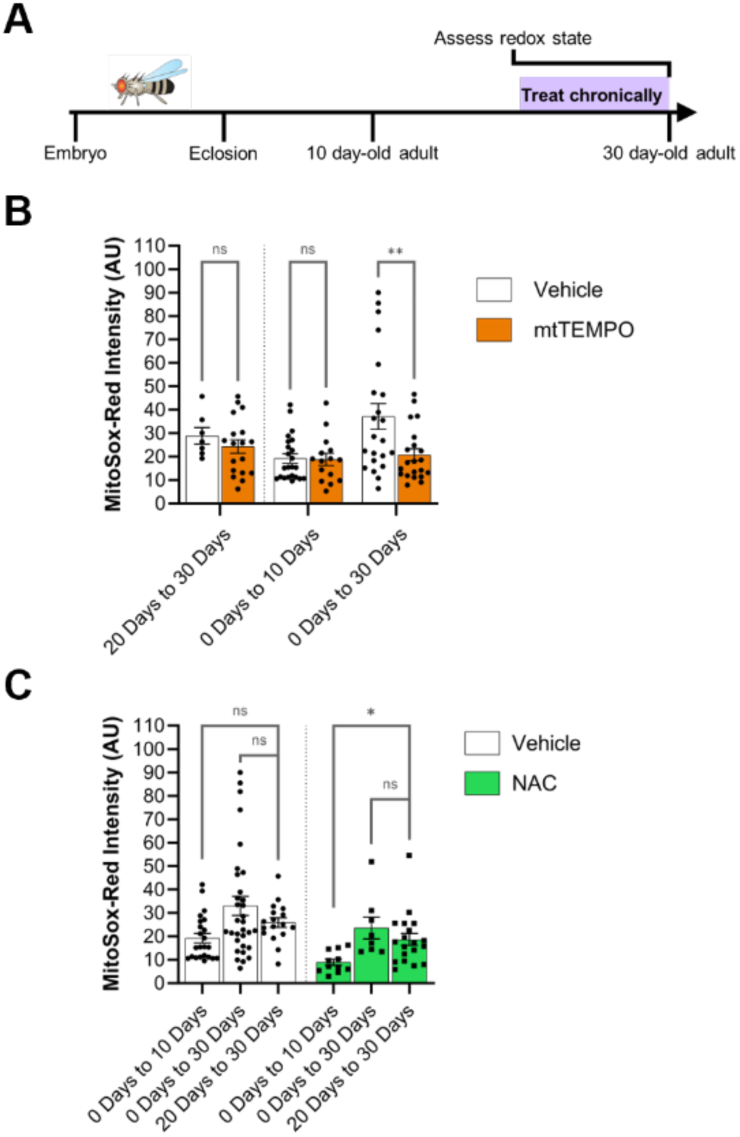
Effects of redox-altering drug treatment duration vs. age at treatment assessment. (A) Schematic of experimental design. WT flies were raised until 20 days old without drug treatment before being fed drugs chronically until 30 days old. The mitochondrial superoxide concentration was then assessed in these flies at 30 days old and compared to data from Figure 3. (B) Comparison of mitochondrial superoxide concentration in flies fed mtTEMPO *vs.* vehicle from 20 days old until 30 days old. Data from quantification of mitochondrial superoxide concentration of flies fed mtTEMPO from 0 days old until either 10 days old or 30 days old is the same as that presented in Figure 3. Vehicle-treated groups were compared to mtTEMPO-treated groups for each chronic feeding paradigm by two-way ANOVA with Šidák’s multiple comparison test. *p* values: * < .05; ** < .01, *** < .001, **** < .0001. (C) Comparison of mitochondrial superoxide concentration among flies fed NAC or among flies fed vehicle from 20 days old until 30 days old. Data from quantification of mitochondrial superoxide concentration of flies fed NAC from 0 days-old until either 10 days old or 30 days old is the same as that presented in Figure 3. Vehicle-treated groups and NAC-treated groups were compared using one-way ANOVAs with Dunnett’s multiple comparison tests. *p* values: * < .05; ** < .01, *** < .001, **** < .0001.

### Comparative transcriptomics of acute vs. chronic redox-altering drug treatment

It is apparent that not only the redox state, but the compensatory metabolic response to a perturbed redox state is important for survival. Thus, we sought to identify the mechanism of this compensatory response. One potential way that compensation could be taking effect is through changes at the level of gene expression. We, therefore, selected two drugs that showed evidence of chronic compensation, 2-AAPA and mitoQ, isolated brains from 10 day-old WT animals that had been treated either acutely or chronically, and performed bulk RNAseq analysis (Fig. 5A). Transcriptomic profiling of these flies revealed small but statistically significant differences in gene expression between the acute and chronic administration conditions, but these were attributable to effects of vehicle treatment (Fig. 5B-C; Supp. Fig. 3). We did not observe changes that correlated with the identity of the specific drug (Fig. 5). This suggests that compensation to these specific chronic drug treatments is likely to occur post-transcriptionally.

**Figure 5.**
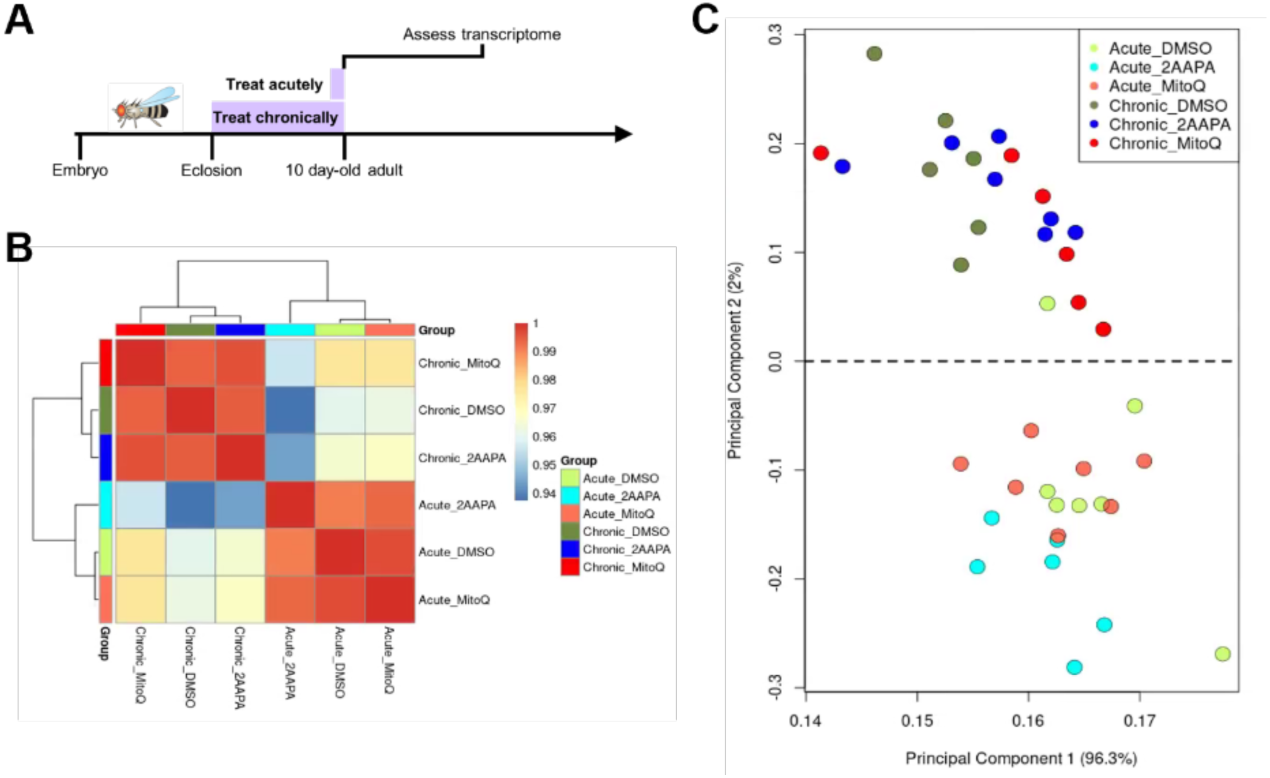
Comparative transcriptomics of acute vs. chronic redox-altering drug treatment. (A) Schematic of experimental design. WT flies were fed drugs either acutely (for 1 day) or chronically (for 10 days) before brains were dissected for transcriptomic analysis at 10 days old. (B) Heatmap of correlation amongst experimental groups based on 184 differentially expressed genes that contribute significantly to the variance amongst samples. Selected genes had a linear fold change of means > |1.5| and a corrected *p* < .05. (C) PCA plot of 184 differentially expressed genes that contribute significantly to the variance amongst samples. Selected genes had a linear fold change of means > |1.5| and a corrected *p* < .05.

## DISCUSSION

Changes in redox state have been linked to aging and to many neurodegenerative conditions (Hou et al., 2019; Liu et al., 2017; Stefanatos & Sanz, 2018). It has been confusing, however, to understand the nature of this linkage. Are the redox changes seen in different conditions responsible for organismal decline in those circumstances, are they homeostatic changes to counteract pathology, or are they simply changes occurring in parallel? Are ROS purely detrimental metabolic by-products, or could they potentially be beneficial molecules? Do aging or pathology affect the ability of the redox metabolic environment to respond to further changes in the redox state? Here, we find that the answers to these questions are nuanced, varying with the reactive oxygen species and the circumstances. Thus, we find evidence from multiple conditions that one classic reactive oxygen species, H_2_O_2_, can be beneficial for survival, while the survival effect of pharmacologically altering a different species, glutathione, is mediated primarily by the compensatory mechanisms induced by those drugs and not by their direct effects on their ROS targets. We also find that the response to treatment with a redox-altering drug, both in terms of the direct redox response and its impact on survival, vary dramatically depending on the age at which the drug is administered, the duration of the treatment, and the genotype of the individual receiving the drug.

To disentangle the influences of genetics and pharmacology on redox state and organismal viability, we proceeded in three steps. First, to define a baseline, we assessed three reactive oxygen species at multiple ages in the brains of WT flies and those with a genetic mutation, Cdk5α-KO, which disrupts the redox state, shortens lifespan, and induces adult-onset neurodegeneration. Second, we identified a set of drugs that alter mitochondrial redox parameters in the WT brain in well-characterized ways when administered acutely. Finally, we addressed whether the alterations in redox state produced by Cdk5α-KO are causal for reduced lifespan by testing whether drugs that tend to reverse Cdk5α-KO-dependent mitochondrial redox changes also tend to improve viability and whether drugs that exacerbate the redox effects of Cdk5α-KO tend to further reduce survival. In this study, we assayed ROS specifically in the brain because Cdk5 is active only in neural tissue, and we focused our attention on the MB as it is a well-documented target of Cdk5 action (Connell-Crowley et al., 2000, 2007; Spurrier et al., 2018; Trunova & Giniger, 2012)

In Cdk5α-KO, we made the unexpected observation that the concentration of mitochondrial H_2_O_2_ actually diminishes with age in a Cdk5-sensitive region of the brain, the MB. To test whether this decrease in mitochondrial H_2_O_2_ concentration is itself detrimental to survival or is a homeostatic response to pathology, we treated Cdk5α-KO flies with either of two drugs, rotenone and mitoQ, that raise the level of mitochondrial H_2_O_2_, and in each case we found that the drug treatment improves viability. This suggests that increased mitochondrial H_2_O_2_ concentration, rather than being detrimental to survival, may actually be beneficial. Further tests reinforce this conclusion as raising the mitochondrial H_2_O_2_ concentration in WT, by chronic treatment with IM, also improves survival. It is interesting to note that the beneficial consequences for survival from increasing mitochondrial H_2_O_2_ concentration in multiple conditions runs counter to the notion that increases in ROS are necessarily detrimental. It has been argued that protective effects of drugs that increase ROS levels are an example of hormesis, wherein a temporary increase of biological stress induces long-term survival benefits, through the activation of genetic programs that counteract the effect (Bárcena et al., 2018; Cheng et al., 2023; Ristow & Zarse, 2010; Tapia, 2006). However, if this were the case, we should observe a long-term reduction in mitochondrial H_2_O_2_ concentration in the rescued condition, whereas we observe an increase. This argues against the simplest version of the hormesis model in this case, though more complex models are possible. The mechanisms by which mitochondrial H_2_O_2_ produces its pro-survival effect are unclear, though one could speculate about potential interactions with redox-sensitive signaling molecules or transcription factors (Lennicke & Cochemé, 2021).

Analyses of the changes in mitochondrial glutathione redox potential provide informative contrasts to the results of mitochondrial H_2_O_2_ concentration. We observed reduction of the mitochondrial glutathione redox potential in the MB of Cdk5α-KO, but drugs that blocked this reduction did not consistently promote survival. For example, NAC prevents the reduction of the mitochondrial glutathione redox potential in Cdk5α-KO relative to WT but did not alter survival of Cdk5α-KO significantly, suggesting that the reduction in mitochondrial glutathione redox potential in Cdk5α-KO is not a major determinant of survival in this genotype. However, 2-AAPA, rotenone, and IM, three drugs that reduce the mitochondrial glutathione redox potential after acute administration, do rescue survival in Cdk5α-KO despite not altering the final mitochondrial glutathione redox potential after chronic administration. Additionally, in the case of chronically treating Cdk5α-KO with 2-AAPA, there are no other changes in mitochondrial redox parameters that might provide an obvious hypothesis for a mechanism. While we cannot rule out the effects of these drugs on other ROS that we haven’t measured, one intriguing possibility is that the mechanisms responsible for the rescue of survival are the compensatory mechanisms that prevent chronic 2-AAPA, rotenone, and IM exposure from affecting the final mitochondrial glutathione redox potential after treatment with these drugs, rather than the direct effect of the drugs themselves.

Our data also do not reveal obvious evidence for detrimental effects of changes in mitochondrial superoxide concentration. It is interesting to note that superoxide is often argued to be a particularly damaging reactive oxygen species, yet we observe an age-related increase in the brains of WT flies that is not associated with any survival effect and suppressing this age-related increase does not impact survival, which argues against the notion that superoxide is detrimental to survival in this context (Shields et al., 2021).

We observe here multiple instances where a redox parameter is changed after acute exposure of a drug but not after chronic exposure of the same drug, suggesting that compensatory metabolic changes occur to alter drug uptake, metabolism or final effect. For example, acute feeding of IM or 2-AAPA to 10 day-old WT flies induces a decrease in mitochondrial superoxide concentration or a reduction in the mitochondrial glutathione redox potential, respectively; however, upon chronic feeding of these drugs to WT flies, these flies show neither the expected reduction in mitochondrial superoxide concentration nor glutathione redox potential. Apparently, the redox state is very malleable after acute administration of redox-altering drugs but less so after chronic administration. The simplest explanation is that there is a compensatory reprogramming of cellular metabolism in response to the drug, whether by altering its bioavailability or its downstream effects. These mechanisms cannot be distinguished unambiguously from the data here, though the occurrence of survival effects in the absence of net changes in redox state upon chronic treatment tends to favor the idea that the effects act downstream. Our transcriptomic analysis of flies fed drugs acutely or chronically indicates that this compensatory reprogramming of metabolism is likely to occur post-transcriptionally, at least for the drugs we have tested, although more investigation is required to identify the mechanisms that are responsible.

It appears that the capacity for compensation to redox-altering drugs is both age- and genotype-dependent. For example, in WT flies, mtTEMPO induces an acute decrease in mitochondrial superoxide concentration, but this effect disappears after chronic exposure for 10 days. However, after 30 days of chronic drug exposure, the effects on mitochondrial superoxide reemerge, suggesting that age and/or treatment duration affects the ability to compensate for the effects of these drugs. Likewise, compensation can be influenced by genotype as we observe that the increase in mitochondrial H_2_O_2_ after acute administration of rotenone and mitoQ is blocked after 10 days of chronic treatment in WT, but not in Cdk5α-KO. Additionally, we observe that compensation can have genotype-dependent effects on survival. For example, the acute effect of 2-AAPA on mitochondrial glutathione redox potential is compensated for (i.e., suppressed) when administered chronically in both genotypes, yet, interestingly, 2-AAPA decreases survival in WT and rescues survival in Cdk5α-KO. Importantly, other changes in redox parameters or compensation cannot account for these survival changes in this case, at least for the ROS we have measured. These differing effects of compensation on survival suggest that there is a difference in the capacity or character of this compensation in the context of a mutation that induces neurodegeneration, and this may partly explain why redox state changes in the context of age-related neurodegeneration are variably correlated with pathology and overall organismal health. Indeed, others have commented on the differing metabolic environments that exist in disease states and how reprogrammed metabolisms can alter responses to pharmacologic manipulations, for example (Parkhitko et al., 2020). It is also intriguing to speculate that metabolic reprogramming after mitochondrial redox-altering drug treatment may influence survival by altering other mitochondrial-based processes besides redox state that have been shown to influence lifespan across numerous species, such as oxidative phosphorylation, beta-oxidation, and the mitochondrial unfolded protein response (Bar-Ziv et al., 2020; Johnson & Stolzing, 2019; Parkhitko et al., 2020; Scialo et al., 2013; Wodrich et al., 2022). It may also be that the observation that both aged flies and Cdk5α-KO flies are less capable of compensation to redox-altering drugs is related to the previous finding that Cdk5α-KO leads to an acceleration of aging (Spurrier et al., 2018). This may suggest that aging and degenerative pathologies that accelerate aging like Cdk5α-KO may be less amenable to treatment with drugs that exert their beneficial effects through a rewiring of metabolism rather than a direct effect on their target. It also implies that caution must be used whenever effects of a drug tested on a young, healthy population are extrapolated to a population that is aged or infirm.

Another unexpected finding from our study was the poor correlation between the predicted effects of a drug and its effects in the brain. This may arise from the use of predictions based on the *in vivo* effects of these drugs in tissues other than the brain or from predictions derived from experiments in simplified systems, such as measurements of ROS obtained from isolated mitochondria that lack the interactions found amongst native cells and tissues (Sanz, 2016). The discordance between the predicted effect of a drug and observed effect in the brain does raise the possibility that the effect of the drug on viability arises due to the effect of the drug in other tissues; however, while we cannot formally rule out this possibility, the effects of these drugs on Cdk5-associated survival are likely occurring due to the effects of the drug in the brain, since this is the tissue where Cdk5 activity is restricted and that is sensitive to Cdk5 pathology. Importantly, this discordance also hints at a potential explanation for why some drugs, for example, antioxidants, fail to deliver beneficial effects in preclinical and clinical trials: the effect of a drug *in vivo* may cause the opposite of its intended effect in the target tissue (Forman & Zhang, 2021; Halliwell, 2024).

The data presented here have implications for both how we think about ROS in disease processes and how we approach ROS as a target for therapeutics. In recent years, it has become clear that the effects of ROS cannot be considered in a monolithic way. Above, we have found that whether altering a reactive oxygen species *in vivo* is beneficial or detrimental depends on the identity of the molecule in question and the age and genotype of the individual. Additionally, whether a particular redox-altering drug could act as a beneficial therapeutic depends on those same variables and also on the age at treatment and the duration of treatment. Furthermore, the net outcome of a drug effect for the individual can be due to the direct effect of the drug and/or the compensatory mechanism(s) it activates. These data underscore the need to separate the many influences of cell and tissue biology on ROS and of ROS on that biology to better understand pathogenesis and develop more effective therapeutics.

## METHODS

### Fly maintenance, stocks, and aging

All flies were housed at 25°C and 50% humidity on a 12:12 hour light:dark cycle. Unless otherwise specified, flies were fed a standard cornmeal-molasses *Drosophila* media (Caltech media; KD Medical, Columbia, MO). All experiments were performed in male adults in an Oregon Red (w^+^) background. The stocks used in these experiments were as follows: *201Y-GAL4* (BDSC, #4440), *UAS-mito-roGFP2-Grx1* (BDSC, #67664), *UAS-mito-roGFP2-Orp1* (BDSC, #67667). The Cdk5α-KO condition (*w^+^; Cdk5α ^20C^/Df(Cdk5α)^C2^; +*) has been described in detail elsewhere (Connell-Crowley et al., 2000, 2007; Spurrier et al., 2018; Trunova & Giniger, 2012).

For the assessment of mitochondrial glutathione redox potential, the MB γ-neuron specific *GAL4* driver *201Y-GAL4* was used to express *UAS-mito-roGFP2-Grx1* in WT and Cdk5α-KO conditions. The resulting experimental genotypes were: WT (*w^+^; 201Y-GAL4/UAS-mito-roGFP2-Grx1; +*) and Cdk5α-KO (*w^+^; Cdk5α^20C^, 201Y-GAL4/Df(Cdk5α)^C2^, UAS-mito-roGFP2-Grx1; +*). For the assessment of mitochondrial H_2_O_2_ concentration, *201Y-GAL4* was used to express *UAS-mito-roGFP2-Orp1* in WT and Cdk5α-KO conditions. The resulting experimental genotypes were: WT (*w^+^; 201Y-GAL4/UAS-mito-roGFP2-Orp1; +*) and Cdk5α- KO (*w^+^; Cdk5α^20C^, 201Y-GAL4/Df(Cdk5α)^C2^, UAS-mito-roGFP2-Orp1; +*). For the all other experiments, the experimental genotypes used were: WT (*w^+^; +; +*) and Cdk5α-KO (*w^+^; Cdk5α ^20C^/Df(Cdk5α)^C2^; +*).

For aging of adult flies, males and females were collected within 24 hours of eclosion and transferred to fresh vials. After three days, males were collected and placed into fresh vials. Flies were aged until they reached the appropriate experimental age. Flies were flipped to fresh vials twice weekly while aging.

### Acute and chronic redox drug feeding

Drug stocks were prepared by dissolving each drug in an appropriate vehicle solvent (Supp. Table 1). Note that there were no within-group differences based on the vehicle used. Drug stocks were aliquoted and stored at −20°C. For acute drug feeding, flies were first maintained and aged on standard cornmeal-molasses *Drosophila* media as described above for the first 9 days after eclosion. Drug stocks were then diluted with H_2_O supplemented with 5% sucrose (w/v) to the experimental concentration. A Kimwipe (Kimberly-Clark, Irving, TX) was placed into an empty vial, and 1 mL of drug or vehicle solution was then added. Flies were then transferred to the vial containing drug or vehicle solution and maintained as described above for 24 hours. For chronic drug feeding, drug stocks were first diluted with H_2_O to the experimental concentration. Drug or vehicle media was then prepared by mixing equal volumes of drug or vehicle solution at the experimental concentration and Formula 4-24 Instant *Drosophila* Medium (Carolina Biological Supply Company, Burlington, NC). Flies were then transferred to drug or vehicle media and maintained and aged as described above with flipping to fresh drug or vehicle media twice weekly while aging.

### General confocal microscopy and image processing

All images were acquired using a LSM 880 confocal microscope (Zeiss, Oberkochen, Germany). All images for a given experiment used identical imaging settings. Note that because of differences in imaging settings between experiments, raw values may differ between experiments. All image analysis was done using ImageJ (version 1.53t, NIH, Bethesda, MD). All image visualizations were done using ImageJ and Imaris software (version 9.5, Oxford Instruments, Abingdon, United Kingdom). Z-stack images were visualized in Imaris using the 3D View tool. Images were rotated and cropped to remove out-of-plane signal as necessary.

### Mitochondrial superoxide concentration assessment

To stain brains with MitoSox-Red, brains were first dissected in 1x PBS before incubating with 30

μM MitoSox-Red (Thermo Fisher Scientific, Washington, DC; #M36008) in 1x PBS for 10 minutes at room temperature protected from light. Brains were then twice washed with 1x PBS for 2 minutes. Brains were mounted on slides between two number 1 glass coverslips (to prevent squishing of brains) and covered with Vectashield mounting medium (Vector Laboratories, Newark, CA) and a coverslip.

Brains were imaged immediately after dissection and incubation using a 10x (0.45 NA) air objective to acquire a Z-stack with a 4.0 µm step size covering approximately 120 µm. The laser settings were excitation at 561 nm and emission between 567-625 nm. To calculate the relative concentration of mitochondrial superoxide, all images were first converted to 32-bit format and thresholded with values below the threshold set to “Not a Number”. A ROI was then drawn around the entire brain, and the total intensity and area of the pixels with numeric values was calculated through the Z-stack. All images were then normalized to the average background fluorescence.

### Cytosolic ROS concentration assessment

To stain brains with H_2_DCFDA, brains were first dissected in 1x PBS before incubating with 5 μM H_2_DCFDA (Thermo Fisher Scientific, Washington, DC; #D399) for 10 minutes at room temperature protected from light. Brains were then thrice washed with 1x PBS for 1 minute. Brains were mounted on slides between two number 1 glass coverslips (to prevent squishing of brains) and covered with Vectashield mounting medium (Vector Laboratories, Newark, CA) and a coverslip.

Brains were imaged immediately after dissection and incubation using a 10x (0.45 NA) air objective to acquire a Z-stack with a 4.0 µm step size covering approximately 120 µm. The laser settings were excitation at 488 nm and emission between 490-560 nm. To calculate the relative concentration of cytosolic ROS, all images were first converted to 32-bit format and thresholded with values below the threshold set to “Not a Number”. A ROI was then drawn around the entire brain, and the total intensity and area of the pixels with numeric values was calculated through the Z-stack.

### Mitochondrial glutathione redox potential and H_2_O_2_ concentration assessment

To determine the dynamic range of the *201Y>mito-roGFP2-Grx1* or *201Y>mito-roGFP2-Orp1* biosensors in the *Drosophila* mushroom body, 10 day-old WT brains were dissected in 1x PBS supplemented with either 2 mM diamide (DA) or 20 mM dithiothreitol (DTT) to oxidize or reduce the biosensors, respectively. Brains were then incubated in DA or DTT for an additional 20 minutes. Brains were then incubated in 30 mM *N-*Ethylmaleimide (NEM) in 1x PBS for 20 minutes at room temperature in order to stabilize the redox-sensitive fluorophore and protect against fixation-induced thiol oxidation (Albrecht et al., 2011). Afterwards, brains were washed in 1x PBS for 5 minutes followed by fixation in 4% paraformaldehyde for 1 hour at room temperature. Brains were then washed thrice in 1x PBS for 10 minutes. Finally, brains were mounted on slides between two number 1 glass coverslips (to prevent squishing of brains) and covered with Vectashield mounting medium (Vector Laboratories, Newark, CA) and a coverslip.

To assess changes in the biosensors in the context of Cdk5α-KO or drug administration, the above procedure was followed with the following changes: brains were dissected in 30 mM NEM in 1x PBS, omitting the dissection and incubation in DA or DTT.

Each MB hemisphere was imaged using a 40X (1.2 NA) water objective to acquire a Z-stack with a 1.0 µm step size covering approximately 120 µm. The laser settings were as follows: 405nm channel (ex: 405nm and em: 500-550nm), 488nm channel (excitation at 488nm and emission between 500-550nm). To calculate the oxidation/reduction ratio of mitochondrial glutathione redox potential and mitochondrial H_2_O_2_ concentration (from *201Y>mito-roGFP2-Grx1* or *201Y>mito-roGFP2-Orp1*, respectively), all images were first converted to 32-bit format and thresholded with values below the threshold set to “Not a Number”. The 405 nm channel was then divided by the 488 nm channel pixel-by-pixel to create a 405nm/488nm ratiometric image. A ROI was then drawn around the mushroom body on this ratiometric image, and the total intensity and area of the pixels with numeric values was calculated through the Z-stack. The reported 405nm/488nm ratio is the average intensity per area of the ratiometric image. Note that this ratiometric measurement corrects for any differences in the absolute expression level of these biosensors.

### Survival assay

To assay survival, flies were chronically fed drug or vehicle media and aged as described above. When flies were flipped to fresh vials, the number of dead and censored male flies were counted. Any fly that escaped the vial, became stuck to the wall or plug of the vial, or became stuck to the food was censored. After 30 days, any flies that remained alive were counted.

### Transcriptomic analysis

For isolation of RNA from 10 day-old WT flies fed drug or vehicle either acutely or chronically, groups of 20 male flies were collected without the use of CO_2_ sedation by flash freezing in liquid nitrogen. Fly heads were recovered by vortexing to separate heads from bodies and subsequently passing the heads and bodies through two sieves (with the upper sieve retaining the bodies and the lower sieve retaining the heads). Groups of heads were then homogenized in Lysis Binding Mix (from MagMAX *mir*Vana Total RNA Isolation Kit; Thermo Fisher Scientific, Washington, DC; #A27828) using Navy Eppendorf RNA Lysis Kits (Next Advance, Troy, NY; #NAVYE5-RNA) and a Bullet Blender Storm 24 (Next Advance, Troy, NY; #4116-BBY24M). RNA was then isolated from these homogenates using the MagMAX *mir*Vana Total RNA Isolation Kit per the manufacturer’s instructions in conjunction with the KingFisher Apex Purification System with 96 Deep-Well head (Thermo Fisher Scientific, Washington, DC; #5400930). The quality of RNA was assessed by High Sensitivity RNA ScreenTape (Agilent Technologies, Santa Clara, CA).

For RNA sequencing, the quality of RNA was first assessed by TapeStation 2200 (Agilent Technologies, Santa Clara, CA) and NanoDrop (Thermo Fisher Scientific, Washington, DC). cDNA libraries were generated from 500 ng of total RNA using NEBNext Ultra II Directional Poly-A RNA Library Prep Kit for Illumina (New England Biolabs, Ipswich, MA; #E7760). cDNA library quality and quantity were assessed by TapeStation 2200 and Qubit (Thermo Fisher Scientific, Washington, DC). cDNA libraries were sequenced on a NextSeq 2000 machine using NextSeq 2000 P3 reagents (Illumina, San Diego, CA).

For analysis of sequencing data, FASTQ files were first quality inspected using the FastQC (available at: https://www.bioinformatics.babraham.ac.uk/projects/fastqc/) and MultiQC (Ewels et al., 2016) tools. For reference mapping (BDGP6), the nf-core RNA-Seq Seq pipeline (version 3.10.1; available at https://nf-co.re/rnaseq) was used in conjunction with select parameters (--outFilterMultimapNmax 500, -- winAnchorMultimapNmax 500, --clip_r1 12, --clip_r2 12). To enumerate counts for known genes, the featureCounts tool was used (Liao et al., 2014). To enumerate counts for transposable elements (TEs), the TEcount tool was used (Jin et al., 2015). These counts were then organized in matrix form, with features in rows and samples in columns. Features observed to not have at least one sample with a value greater than zero were filter removed. This filtered matrix was then imported into R, the matrix of counts was pedestalled by 2, log2 transformed, then cross-sample normalized. For cross-sample normalization, cyclic LOESS was applied using the normalizeBetweenArrays function available as part of the limma package. Post normalization, outliers were detected by covariance-based principal component analysis scatterplot and removed. Cross-sample normalization was repeated using transformed counts for surviving samples then noised modeled by experiment condition (CV∼mean) using the LOWESS function. Resulting fits were visually inspected and the noise cut-off for the data defined to equal the lowest mean value where the linear relationship between CV and mean was grossly lost (1.75). Features not having at least one sample value greater than the noise cut-off value were discarded with any value less floored to equal the noise cut-off value. Features were also discarded if they were observed to have a CV greater than the average CV observed at the noise cut-off value (80%). To identify differential expressed features for each possible pairwise comparison of experiment conditions, the ANOVA test under Benjamini-Hochberg, false discovery rate, and multiple comparison correction conditions was used followed by post-hoc testing using the TukeyHSD test. Features observed to have an ANOVA corrected *p* < .05, a post-hoc *p* < .05, and an absolute linear fold change of means > |1.5| for a comparison were deemed to be differentially expressed for the comparison, respectively. Post testing, sample-to-sample relationships were inspected by covariance-based PCA scatterplot and correlation-based heatmap using the union set of differential features across all comparisons. Differential analysis was then repeated using ANCOVA in place of ANOVA and general linear hypothesis testing in place of TukeyHSD while adjusting for drug (DMSO, 2AAPA, MitoQ) in one pass and adjusting for drug schedule (acute, chronic) in a second, separate pass.

All sequencing data will be made available on GEO prior to publication.

### Statistical analyses

All statistical analyses were performed using Prism (GraphPad, version 10, Boston, MA). Specific statistical tests are specified in the Figure Legends. Data were tested for normality using the Kolmogorov-Smirnov test and for equality of variances using the F-test of equality of variances or Bartlett’s test. Outliers were detected using the ROUT method with Q = 1% (Motulsky & Brown, 2006). All analyses were completed blind to genotype and age. One biological replicate is represented by a single fly, unless otherwise specified. When appropriate, measurements from each MB hemisphere from a single brain were treated as technical replicates, and these measurements were averaged together. All graphs show mean +/- SEM unless otherwise noted in the Figure Legends.

## Supporting information

Supplemental datasheet

## ACKNOWLEDGEMENTS

We wish to thank all current and former members of our lab for helpful discussions, comments, and insights. Specifically, we are thankful to Arvind Shukla, Andrew Scott, Rameen Forghani, Abigail Molnar, Phil McQueen, Hitesh Chaouhan, and Suparna Saha. We are also thankful to Joy Gu and Irina Kuzina for technical assistance. We are also thankful to Bill Rebeck, Tom Coate, Derek Narendra, and Tingting Wang for helpful insights, discussions, and suggestions during the course of these experiments and for comments on the manuscript. We are also grateful for the microscopy assistance of Stephen Wincovitch of the National Human Genome Research Institute (NHGRI) Cytogenetics and Microscopy Core Facility. We are also thankful to Abdel Elkahloun of the NHGRI for his invaluable support with RNA sequencing. We also thank Kory Johnson of the National Institute of Neurological Disorders and Stroke (NINDS) Bioinformatics Core for assistance with the RNA sequencing bioinformatics. We are also thankful to Derek Narendra of the NINDS for sharing lab space and equipment as well as Mark Cookson of the National Institute on Aging (NIA) for sharing equipment. This work was supported by the Basic Neuroscience Program of the Intramural Research Program of the NINDS / National Institutes of Health (NIH) (Z01 NS003106 to E.G.). All authors report no conflicts of interest. All relevant data can be found within the article or in the supplementary information (Supp. Table 3).

## AUTHOR CONTRIBUTIONS

APKW: conceptualization, methodology, validation, formal analysis, investigation, writing – original draft, writing – review & editing, visualization

BTH: writing – review & editing, supervision

EG: conceptualization, resources, writing – review & editing, supervision, funding acquisition

## CONFLICT OF INTEREST STATEMENT

The authors declare no conflicts of interest.

## ABBREVIATIONS

3-AT: 3-amino-1,2,4-triazole
BSO: buthionine sulfoximine
Cdk5: cyclin dependent kinase 5
Cdk5-KO / KO: knockout of Cdk5α
DDC: diethyldithiocarbamate
IM: imisopasem manganese
MB: mushroom body
mitoQ: mitoquinone
mtTEMPO: mito-TEMPO
NAC: N-acetyl cysteine
PQ: paraquat
ROS: reactive oxygen species
WT: wild-type

## SUPPLEMENTAL FIGURES AND TABLES

**Supplementary Figure 1.**
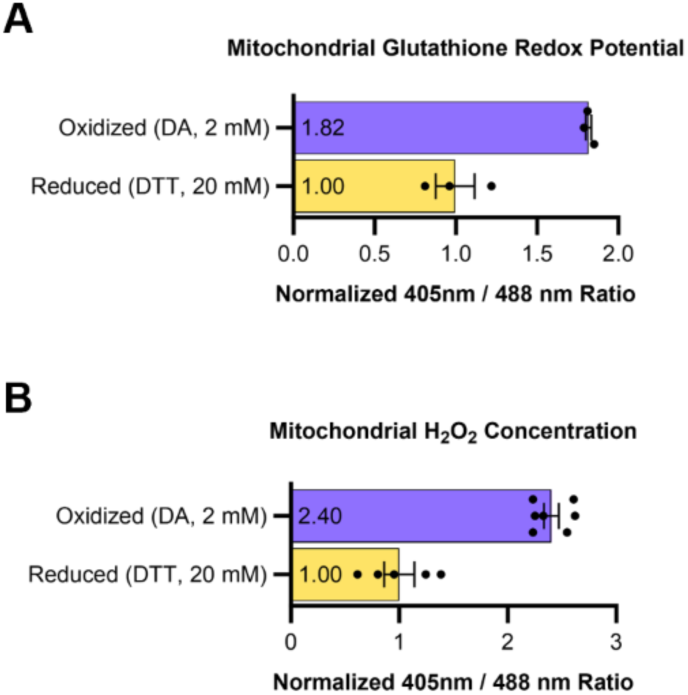
mito-roGFP2 biosensors capture redox changes in the *Drosophila* mushroom body. (A-B) Oxidation (with DA) or reduction (with DTT) of 10 day-old WT flies expressing *201Y>mito-roGFP2-Grx1* (A) or *201Y>mito-roGFP2-Orp1* (B) measuring the mitochondrial glutathione redox potential or the H_2_O_2_ concentration, respectively. 405nm/488nm ratios have been normalized by setting the average reduced 405nm/488nm ratio to 1. A higher 405nm/488nm ratio indicates a more oxidized mitochondrial glutathione redox potential (A) or a higher mitochondrial H_2_O_2_ concentration (B).

**Supplementary Figure 2.**
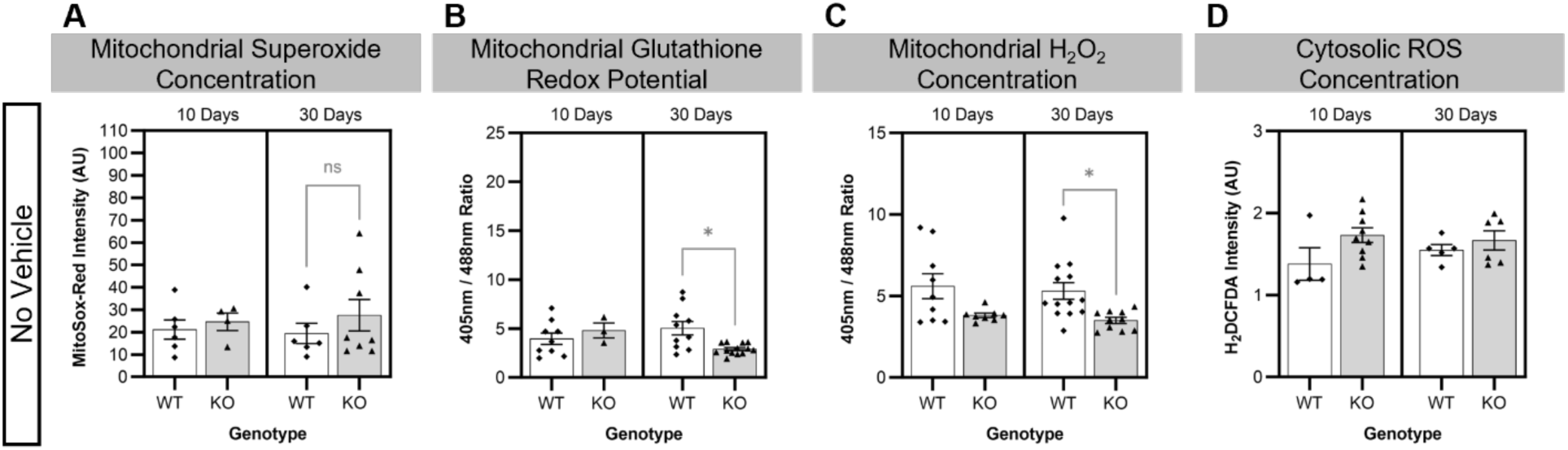
WT and Cdk5α-KO flies raised on Caltech media and not exposed to vehicle show similar changes in redox parameters to flies exposed to vehicle. (A) Quantification of mitochondrial superoxide concentration in WT or Cdk5α-KO brains. A higher MitoSox-Red intensity indicates a higher mitochondrial superoxide concentration. *p* values: * < .05; ** < .01, *** < .001, **** < .0001. (B) Quantification of mitochondrial glutathione redox potential in WT or Cdk5α-KO MBs. A larger 405nm/488nm ratio indicates a more oxidized mitochondrial glutathione redox potential. *p* values: * < .05; ** < .01, *** < .001, **** < .0001. (B) Quantification of mitochondrial H_2_O_2_ concentration in WT or Cdk5α-KO MBs. A larger 405nm/488nm ratio indicates a higher mitochondrial H_2_O_2_ concentration. *p* values: * < .05; ** < .01, *** < .001, **** < .0001. (D) Quantification of normalized H_2_DCFDA in WT or Cdk5α-KO brains. A higher H2DCFDA intensity indicates a higher cytosolic ROS concentration. *p* values: * < .05; ** < .01, *** < .001, **** < .0001.

**Supplementary Figure 3.**
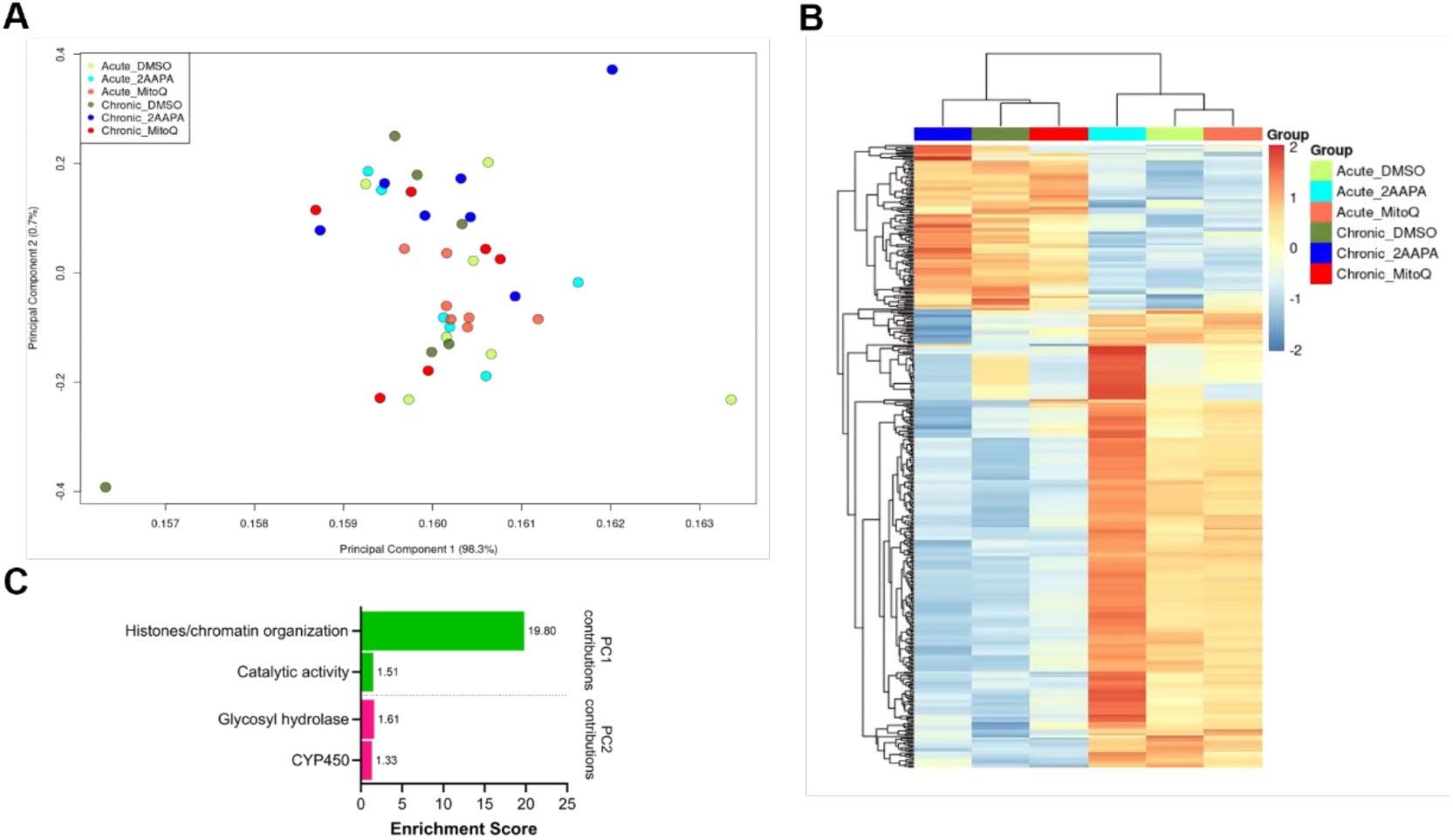
Transcriptomic profiling of acute and chronically fed 10 day-old WT flies reveals changes associated with vehicle treatment between acute and chronic groups, but not among drug treatments. (A) PCA plot of 10 day-old WT flies fed either DMSO (vehicle), 2-AAPA, or mitoQ acutely or chronically. (B) Bar graph showing the enrichment score of the functional annotation clusters calculated by DAVID 6.8 of 184 differentially expressed genes that contribute significantly to PC1 and PC2 (Huang et al., 2009; Sherman et al., 2022). (C) Gene heatmap of 184 differentially expressed genes that contribute significantly to PC1 and PC2. Selected genes had a linear fold change of means > |1.5| and a corrected *p* < .05.

**Supplementary Table 1.** Summary of drugs used in this study. Drugs used in this study with CAS identification number, supplier information, solvents and concentrations used to prepare drug for feeding. For non-standard drug abbreviations, a full drug name is included if one exists.

| Drug Abbreviation | Drug Name | CAS Number | Supplier | Catalog Number | Vehicle Solvent | Drug Concentration |
| --- | --- | --- | --- | --- | --- | --- |
| 2-AAPA | | 1133387-90-2 | Sigma-Aldrich | A4111 | DMSO | 10 μM |
| 3-AT | 3-amino-1,2,4-triazole | 61-82-5 | Sigma-Aldrich | A8056 | H<sub>2</sub>O | 500 μM |
| Antimycin A | | 1397-94-0 | Sigma-Aldrich | A8674 | EtOH | 9.11 μM |
| Auranofin | | 34031-32-8 | Sigma-Aldrich | A6733 | DMSO | 1 μM |
| BSO | Buthionine sulfoximine | 83730-53-4 | Cayman Chemical | 14484 | H<sub>2</sub>O | 10 mM |
| DDC | Diethyldithiocarbamate | 20624-25-3 | Sigma-Aldrich | D3506 | H<sub>2</sub>O | 1 mM |
| EUK-8 | | 53177-12-1 | Sigma-Aldrich | SML0742 | DMSO | 500 μM |
| Idebenone | | 58186-27-9 | Sigma-Aldrich | I5659 | DMSO | 10 μM |
| IM | Imisopasem manganese | 218791-21-0 | MedChemExpress | HY-13336 | DMSO | 500 μM |
| mitoQ | Mitquinone mesylate | 845959-50-4 | Cayman Chemical | 29317 | EtOH | 100 μM |
| mtTEMPO | Mito-TEMPO | 1334850-99-5 | Sigma-Aldrich | SML0737 | EtOH | 1 mM |
| NAC | N-acetyl cysteine | 616-91-1 | Sigma-Aldrich | A7250 | DMSO | 1 mM |
| PQ | Paraquat | 1910-42-5 | Sigma-Aldrich | 36541 | H<sub>2</sub>O | 100 μM |
| Rotenone | | 83-79-4 | Sigma-Aldrich | R8875 | DMSO | 10 μM |
| S1QEL1.1 |  | 897613-29-5 | Cayman Chemical | 20982 | DMSO | 800 nM |
| S3QEL2 |  | 890888-12-7 | Cayman Chemical | 18556 | DMSO | 800 nM |

**Supplementary Table 2.** Summary of drugs used in acute drug administration screen and their predicted effects. Drugs used in acute drug administration screen with their mechanisms of action and their predicted effects on survival, mitochondrial superoxide concentration, mitochondrial glutathione redox potential, and mitochondrial H_2_O_2_ concentration. The predicted drug effects are based on their mechanisms of action as well as experimental evidence in various *in vitro* and *in vivo* contexts. The predicted drug effects are indicated by the green up arrow (drug predicted to improve survival), the magenta down arrow (drug predicted to reduce survival), the orange up arrow (drug predicted to lead to an increased oxidation state), the blue down arrow (drug predicted to lead to a decreased oxidation state), or a black bar (drug predicted to led to no change in survival or oxidation state or a prediction cannot be made for the given drug).

| Drug | Target of Action | Effect on Target | Survival | Superoxide | Glutathione | H<sub>2</sub>O<sub>2</sub> |
| --- | --- | --- | --- | --- | --- | --- |
| 2-AAPA | TrxR & GR | Inhibitor | ↓ | — | ↑ | — |
| 3-AT | Catalase | Inhibitor | ↓ | — | — | ↑ |
| Antimycin A | ETC complex III | Inhibitor | ↓ | ↑ | ↑ | ↑ |
| Auranofin | TrxR | Inhibitor | ↓ | — | ↑ | — |
| BSO | GCL | Inhibitor | ↓ | — | ↓ | — |
| DDC | Cu/Zn-SOD | Inhibitor | ↓ | ↑ | ↓ | ↓ |
| EUK 8 | Mn-SOD & catalase | Mimetic | ↑ | ↓ | — | ↑ |
| Idebenone | CoQ(10) | Mimetic | ↑ | ↓ | — | ↓ |
| IM | Mn-SOD | Mimetic | ↑ | ↓ | — | ↑ |
| mitoQ | CoQ(10) | Mimetic | ↑ | ↓ | — | ↓ |
| mtTEMPO | Free radicals | Scavenger | ↑ | ↓ | — | — |
| NAC | Glutathione | Precursor | ↑ | — | — | ↑ |
| Paraquat | Free radicals | Toxicant | ↓ | ↑ | ↑ | ↑ |
| Rotenone | ETC complex I | Inhibitor | ↓ | ↑ | ↑ | ↑ |
| S1QEL1.1 | ETC complex I | Suppressor of electron leak | ↑ | ↓ | — | ↓ |
| S3QEL2 | ETC complex III | Suppressor of electron leak | ↑ | ↓ | — | ↓ |

**Supplementary Table 2. Summary of drugs used in acute drug administration screen and their predicted effects.**
| Key |  |
| --- | --- |
| ↑ | Improves survival |
| ↓ | Reduces survival |
| ↑ | Increases oxidation state |
| ↓ | Decreases oxidation state |
| — | No change |

**Supplementary Table 3.** Data presented in this study. See “Wodrich et al. – Supplementary Data.xlsx” Summary data presented in this study, excluding raw RNA sequencing data (see above).

## Notes

### Competing Interest Statement

The authors have declared no competing interest.

